# Analysis of the testicle’s transcriptome of the Chagas disease vector *Rhodnius prolixus*

**DOI:** 10.1101/616193

**Authors:** Jovino C. Cardoso, Jose M. C. Ribeiro, Daniela V. dos Santos, Marcos H. Pereira, Ricardo N. Araújo, Nelder F. Gontijo, Grasielle C. D. Pessoa, Mauricio R. V. Sant’Anna, Marcos H. F. Sorgine, David Majerowicz, Marcelo Medeiros, Gloria R. C. Braz, Rafael D. Mesquita, Pedro L. Oliveira, Leonardo B. Koerich

## Abstract

*Rhodnius prolixus* is amongst the most important vectors of *Trypanosoma cruzi* in the Americas, putting thousands of people at risk of contracting Chagas Disease. This insect is also one of the most important models in insect physiology, especially regarding the blood-feeding process. However, studies on *R. prolixus* genetics lagged, and our understanding on the regulation of gene expression is incipient. Transcriptomes have the power to study the expression of thousands of genes in a single experiment. A comprehensive *R. prolixus* transcriptome was performed in 2014, sequencing RNA from different tissues (anterior gut, midgut, posterior gut, rectum, ovaries, fat body, maphigian tubules, and testicles). However, on that occasion, only the gut transcriptome was deeply analysed. Here we evaluated the results of the testicles transcriptome of *R. prolixus* with the objective to find and understand genes that could have an important role in male reproduction. We found, that from the 25,673 transcripts assembled in the whole transcriptome, 5,365 have a testicle specific expression pattern. As expected, amongst the most abundant families of transcripts, are those related to spermatogenesis and male fertility, such as myosins, actins, and dyneins. To our surprise, lipocalins, serine protease inhibitors (serpins), and lysozymes also were highly abundant in testicles. The role of these classes of genes are well known in other tissues, such as salivary glands and gut, but very little is known on their role in male reproduction (and we proposed here a few hypothesis that could be tested to address the role of these genes in male fertility). It would be interesting to study further the role of these genes on *R. prolixus* male fertility. Finally, as a reflection of the lack of knowledge on triatomine genetics, we found that almost half of the transcripts in *R. prolixus* testicles have no similarities to any other genes on reference databases. Our study shows that we still have a lot to know and to understand about reproduction in triatomine, especially in males. Besides the large number of genes without described function (possibly novel genes), there are those in which the function is known for other tissues, and we can only guess, at best, the role and importance of such genes for triatomine male fertility.

**Author Summary:** The understanding of the biology of insect’s vectors of parasitic diseases is key to the development of strategies of public health. For decades, the studies on the biology of male insects’ vectors of diseases was neglected, since in many cases female insects are those with relevant role in the spread of diseases. With the development of genomics, large scale studies to compare differential gene expression (transcriptomics) among different tissues, developmental stages, and sex became accessible. In this study, we looked at the physiology of the male reproductive organs of the vector of Chagas disease *Rhodnius prolixus*. This is a first glimpse, from a perspective of genes differentially expressed in male gonads, in such insects. We also performed an effort to link all identified genes with the insect genome published in 2015. We found ~14,000 genes expressed in the testicles, from which 5,635 genes are expressed exclusively in male reproductive organs. From the ~14,000 genes, we were able to attribute putative biological functions to 6,372 genes, which allowed us to draw a bigger picture on how these genes contribute to male fertility. This study now opens the door for further in-depth studies to find key genes for *R. prolixus* reproductive biology.

## Introduction

*Rhodnius prolixus* (Hemiptera:Reduviidae) is the main vector of Chagas disease in Central and the northern part of South America. Chagas disease is amongst the most important parasitic infection in Latin America, and nearly 6 million people are infected with *Trypanosoma cruzi* (the causative agent of Chagas disease) in 21 countries, with ~40,000 new cases every year^1^. In addition, since the works of Sir Vincent Wigglesworth in the first half of the 20^th^ century, *R. prolixus* became a model for the study of insect physiology^2^, especially in respect to hematophagy. However, as in many other hematophagous insects, most studies on *R. prolixus* focused on female biology, while the knowledge on male biology is very incipient.

In recent years, RNA sequencing techniques have become cheaper and easier to access for the scientific community. Whole mRNA sequencing (or whole RNA sequencing in some cases) is now widely used as a tool to discover new genes and new transcript isoforms, to study genes differentially expressed in different conditions (or tissues and developmental stages), and to create a catalogue of gene candidates for further functional studies. As a tradition in vector biology studies, a few transcriptomes have catalogued expression of *R. prolixus* genes in the salivary glands^3^, gut^4^, and ovaries^5^ in order to study hematophagy, vector-host interactions, vector-parasite interactions, and female fertility. In ticks, blood-feeding also stimulates spermatogenesis, mating, and production of male factors that trigger female reproduction^6^. Studies performed in *A. gambiae* have highlighted a profound systemic changes in gene expression in female mosquitoes upon fertilization, due to transfer of material produced by the male sexual apparatus, including ecdysone, with implications on reproductive physiology and vector competence^7^. *R. prolixus* testicles start to develop in fifth instar nymphs and are fully functional in adults. A male adult has two testicles, and each testicle has a certain composition: two long testicles; five short testicles; a vas deferens duct; vesicular seminal duct; duct of accessory glands; and four accessory glands (three transparent and one opaque)^8^. Spermatogenesis occur in each of the seven testicles arms, and we still do not know if the different arms sizes have any specific role on male reproduction. On the other hand, many studies on *R. prolixus* accessory glands have been conducted since the beginning of the 20^th^ century, showing its importance on spermatophore production and male fertility^9^. In insects, the accessory glands are involved in^10^: *i)* facilitation of the insemination of the females; *ii)* sperm activation; *iii)* formation of mating plugs; *iv)* modification of female mating behaviour; *v)* sperm competition; *vi)* and egg maturation and oviposition. Still, little is known about key genes related to male fertility and reproduction biology in insects.

Transcriptomics and proteomics have been successfully applied to create a catalogue of testicle specific genes in arthropods. The testicles transcriptome of *Drosophila melanogaster* described ~8,500 genes and further analyses on gene expression revealed that 399 genes were upregulated in testicles^11^. Most of the upregulated genes are typical components of sperm structure (*eg.* dynein, mitochondria, and outer dense fibre), sugar metabolism, and peptidases. Among those, only dynein was previously known to cause infertility in males. In the leishmaniasis vector *Lutzomyia longipalpis* (Diptera:Psychodidae), the testicles transcriptome suggested that many of the accessory gland proteins are involved in proteolysis (*e.g.* serine protease, metalloproteases, and protease inhibitors), in immunity, and redox metabolism^12^. In the tick *Dermacentor variabilis*, the testicles transcriptome and spermatophore proteome suggested the importance of serine/threonine kinase, metalloendoproteinases, ferritins, serine proteases, trypsin, cysteine proteases, serpins, a cystatin, GPCR, and others, in the reproductive biology of these arthropods^6^.

On the other hand, the physiology of *R. prolixus* male reproduction and its impact on female development are essentially a black box. Consequently, a new global approach using transcriptomics, proteomics, and quantitative gene expression is needed to understand male physiology and fertility of this insect. In 2010, a comprehensive RNA-seq from different tissues (anterior gut, midgut, posterior gut, rectum, Malpighian tubules, ovaries, fat body, and testicles) of *R. prolixus* was performed, and the analysis of the gut transcriptome was published in 2014^4^. In this study, we analysed the data from the testicles transcriptome and found ~14,000 *R. prolixus* transcripts, with 5,365 testicles specific transcripts. This is the first comprehensive genetic study on *R. prolixus* male reproductive organs, and we expect to provide some insight on male specific fertility genes and candidates for further functional studies on this insect reproductive biology.

## Material and Methods

### Insects

Insects used for transcriptome were *Rhodnius prolixus* from a colony kept at UFRJ (Rio de Janeiro), fed with rabbit blood, and raised at 28°C and 70% relative humidity. Adult females (five from each condition) receiving their second blood meal after the imaginal molt were dissected before feeding, twelve hours after blood meal (ABM), twenty-four hours ABM, two days ABM, and five days ABM. A group of males (blood fed, five days ABM) was dissected to obtain the testicles. Organs (anterior midgut, posterior midgut, fat body, ovary, Malpighian tubules, and testicles) were dissected, homogenized in TriZol, and processed as described below. All the animal work was conducted according to the guidelines of the institutional care and use committee (Committee for Evaluation of Animal Use for Research from the Federal University of Rio de Janeiro), which was adapted from the National Institutes of Health Guide for the Care and Use of Laboratory Animals (ISBN 0-309-05377-3). The protocols received registry # 115/13 from the Animal Ethics Committee (Comissão de Ética no Uso de Animais, CEUA).

### RNA extraction, library preparation, and sequencing

Organs were homogenized in TriZol, and total RNA was isolated, followed by mRNA purification using the Micro-Fasttrack 2.0 kit from Invitrogen (San Diego, CA, USA) according to manufacturer’s instructions. Libraries were constructed using the Smart cDNA Library Construction kit from Clontech (Palo Alto, CA, USA) and normalized using the Trimmer cDNA Normalization kit from Evrogen (Moscow, Russia). The libraries were sequenced on a 454 genome sequencer FLX Titanium machine (Roche, Roche 454 Life Sciences, Branford, CT, USA).

### Bioinformatics

Detailed bioinformatic analysis of our pipeline can be found in our previous publication^13^. Pyrosequencing reads were removed from vector and primer sequences by running VecScreen. The resulting assemblies, plus the clean pyrosequenced data, were joined by an iterative BLAST and cap3 assembler. This assembler tracks all reads used for each contig, allowing deconvolution of the number of reads used from each library for tissue expression comparisons using traditional reads per kilobase million (RPKM). Non-testicles RPKM is the sum of reads from all libraries except for the testicle’s library.

Coding sequences were extracted using an automated pipeline based on similarities to known proteins or by obtaining CDS from the larger open reading frame of the contigs containing a signal peptide. A non-redundant set of the coding and their protein sequences was mapped into a hyperlinked Excel spreadsheet, which is presented as S1 Excel Table. Signal peptide, transmembrane domains, cleavage sites, and mucin-type glycosylation were determined with software from the Centre for Biological Sequence Analysis (Technical University of Denmark, Lyngby, Denmark). To assign coding sequences as being of bacterial, viral, or invertebrate origins, the top blastp scores of the deducted proteins against each database were compared. If the ratio between the top two scores was larger than 1.25 and the e value of the blastp against pathogen or vertebrate was smaller than 1e-15, then the CDS was assigned to the top-scoring organism group.

For genome and proteome mapping, transcripts were aligned to the reference genome^13^ (version RproC3) and the reference proteome (version RproC3.3) downloaded from VectorBase. Transcript alignment with genome and proteome was performed using local blastn and blastx, respectively, with the following parameters: word size of 20 for blastn and 3 for blastx; e-value 0.001. We considered a successful alignment to the genome those hits that aligned at least 80% of the transcript sequence with identity above 98%, and a successful alignment to the proteome those hits that aligned at any amount of amino-acids of the transcript sequence to the proteome with identity above 98%.

Raw sequences were deposited on the Sequence Read Archive (SRA) from the NCBI under bioproject accession PRJNA191820. The individual run files received accession numbers SRR206936, SRR206937, SRR206938, SRR206946, SRR206947, SRR206948, SRR206952, SRR206983, and SRR206984. A total of 2,475 coding sequences and their translations were submitted to the Transcriptome Shotgun Assembly (TSA) project deposited at DDBJ/EMBL/GenBank under the accessions GAHY01000001-2475.

### Differential expression analysis

For differential expression analysis, RPKM values were transformed into z-scores, using data from all libraries to calculate the mean Log. For heatmap analysis, we calculated the mean z-score for each protein class based on their putative role (*e.g.* protein synthesis machinery, amino-acid metabolism, detoxification, etc). Heatmap of protein classes was constructed using Heatmapper^45^. Since we were interested only in having an insight in contig abundancy, we did not perform an in-depth differential expression analysis using common approaches (*e.g.* DEGseq or edgeR). Thus, we do not present and discuss any DE contigs in terms of statistical significance.

### Evolutionary analysis

Protein sequences from other organisms were obtained at NCBI and aligned with Muscle^46^. Evolutionary analyses were conducted in MEGA7^47^. The evolutionary history was inferred using the Neighbour-Joining method (10.000 replicates; pairwise deletion)^48^. The evolutionary distances were computed using the Poisson correction method and are in the units of the number of amino acid substitutions per site. All accession numbers are shown in the respective figures.

## Results and Discussion

The original transcriptome^4^ generated a total of 171,124 (454) reads that where assembled in 25,673 contigs. In this work, we mapped all transcripts from the original transcriptome to the *R. prolixus* genome^13^. We were able to map 22,052 (85.9%) transcripts to the *R. prolixus* genome, but only 9,311 (36.3%) were mapped to the insect proteome (S1 Table). We will discuss the mapping results in a specific section by the end of this manuscript.

Regarding differential expression, we found that 14,454 are expressed in testicles, from which 5,365 transcripts have a specific expression pattern (assembled exclusively with reads from testicles library), while 9,089 are transcribed in testicles and in other tissues (Figure 1). We did not observe testicles expression for 11,219 transcripts. Transcript function was predicted by using Blast against the NR, SwissProt, COG, KOG, CDD, and Pfam databases. A total of 7,691 transcripts did not show any similarity with described proteins, possibly representing novel *R. prolixus* genes. From those transcripts showing blast hits, we classified them in seven major categories: housekeeping, secreted, immunity, transposable elements, viral, unknown conserved (proteins with unknown function but found in other organisms), and unknown (S1 Table). Each transcript was then categorized in functional classes (*e.g.* protein synthesis, transcription machinery, signal transduction, etc) or protein families (*e*.g. lipocalins, serine proteases, lysozymes, etc.). Nearly 85% of the testicles contigs belong to unknown (6,395 contigs) and housekeeping (6,012 contigs) categories (Figure 2), with the relative abundance (RPKM) of transcripts following the same pattern. On the other hand, the abundance of unknown transcripts in other tissues is ~2.5-fold higher than the abundance of housekeeping transcripts. Another difference is observed in the secreted transcripts category. While the number of contigs in this category is almost the same for testicles and other tissues (182 and 214 contigs, respectively), the relative abundance of testicles secreted transcripts is ~2-fold higher than in other tissues (Figure 2), suggesting an important role of testicles as a secretion organ.

**Figure 1.**
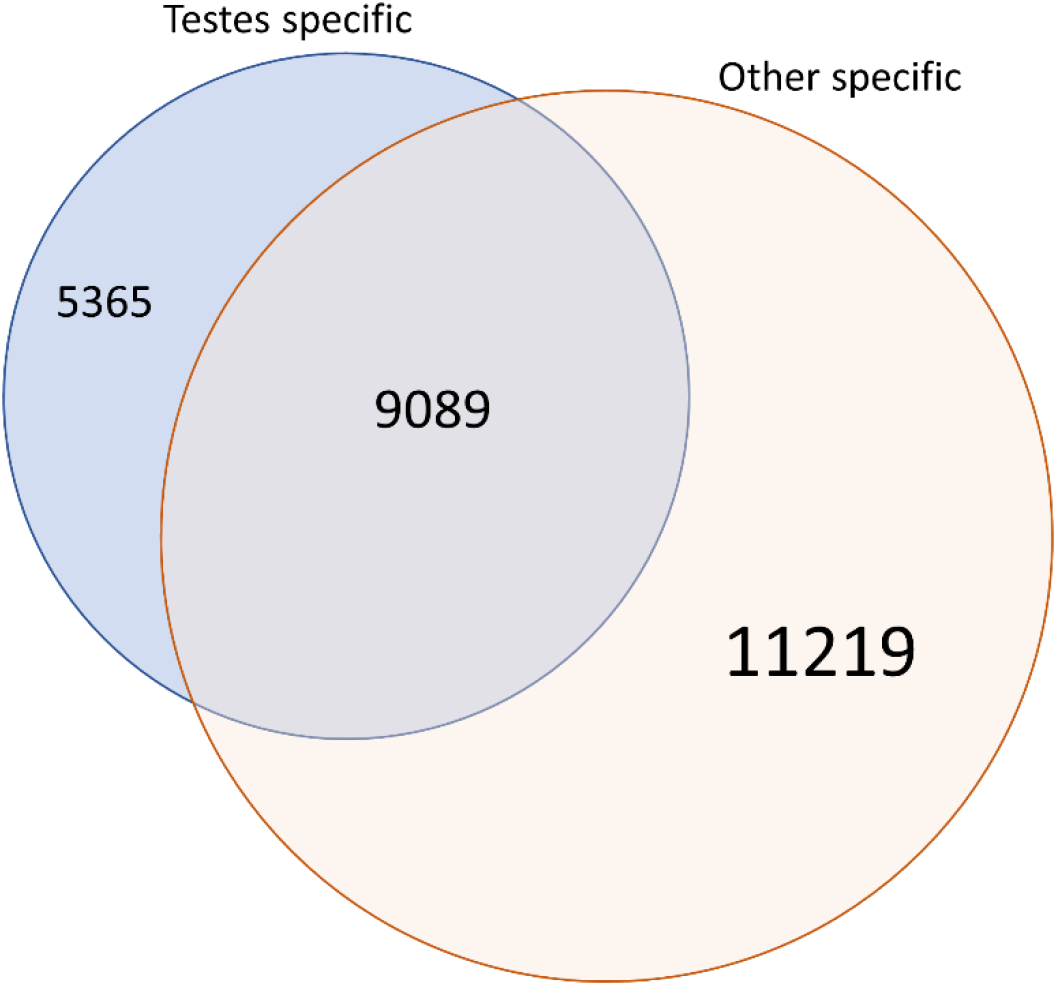
Distribution of specific and non-specific testicles contigs in *Rhodnius prolixus*. Other specific includes transcripts found in gut, ovaries, fat body, and Malpighian tubules.

**Figure 2.**
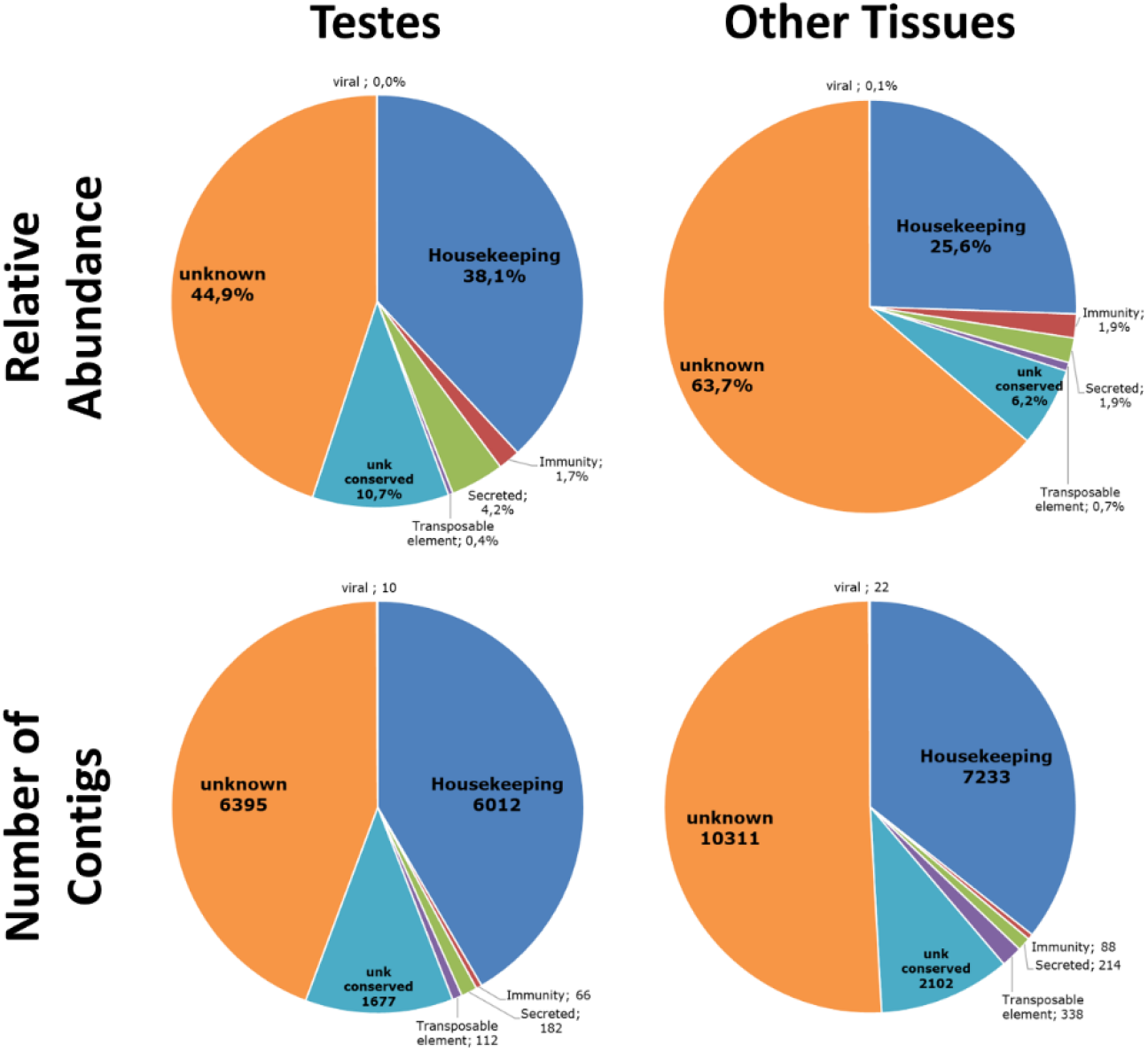
Relative abundance of transcripts and number of contigs by functional class in testicles and other tissues of *Rhodnius prolixus.* Other tissues of *R. prolixus* are gut (anterior gut, midgut, posterior gut, and rectum), ovaries, malpighian tubules. and fat body. Abundance is based in reads per kilobase million (RPKM).

### Overview in housekeeping, secreted and immunity-related genes

#### Housekeeping

When we evaluate the relative abundance of transcripts by functional class in each major category in comparison to other tissues, we observe differences in only a few groups (Figure 3). In the housekeeping category, we observe that there is a trade-off between protein synthesis, cytoskeletal, and signal transduction transcripts (Figure 3 and S2 Table). While the abundance of protein synthesis transcripts decreases ~1.5-fold in testicles in comparison to non-testicles, the abundance of cytoskeletal and signal transduction transcripts increases ~2.5 and ~1.5-fold, respectively. The increase in cytoskeletal transcripts should be expected due to genes involved spermatogenesis and the flagellar structure in spermatozoa, which involves actins, myosins, tubulins and dyneins. Actins and myosins have a key role in mitosis and meiosis, and we found many actin and myosin transcripts overexpressed in testicles. In fact, two of the most highly expressed genes in testicles (Table 1) belong to the myosin family; a myosin class II transcript (1.5-fold higher in testicles) representing ~2% of all testicles transcripts, and a stretchin-mlck, which has expression ~24-fold higher in testicles (representing 0.75% of all testicles transcripts). Tubulins and dyneins are the main components flagella, and we found 38 tubulins and 27 dyneins expressed in testicles. Tubulins represents 7.3% of all testicles’ cytoskeletal transcripts with a RPKM ~3-fold higher in testicles than other tissues, while dyneins represent 4.1% of all testicles’ cytoskeletal transcripts with a RPKM ~20-fold higher in testicles. We also observed discrete abundancies of kinesins (~5-fold increase in testicles in comparison with other tissues) corresponding to 2.9% of all testicles’ cytoskeletal transcripts. Discussion on myosin’s, actins, tubulins, and dyneins are detailed further.

**Table 1.**
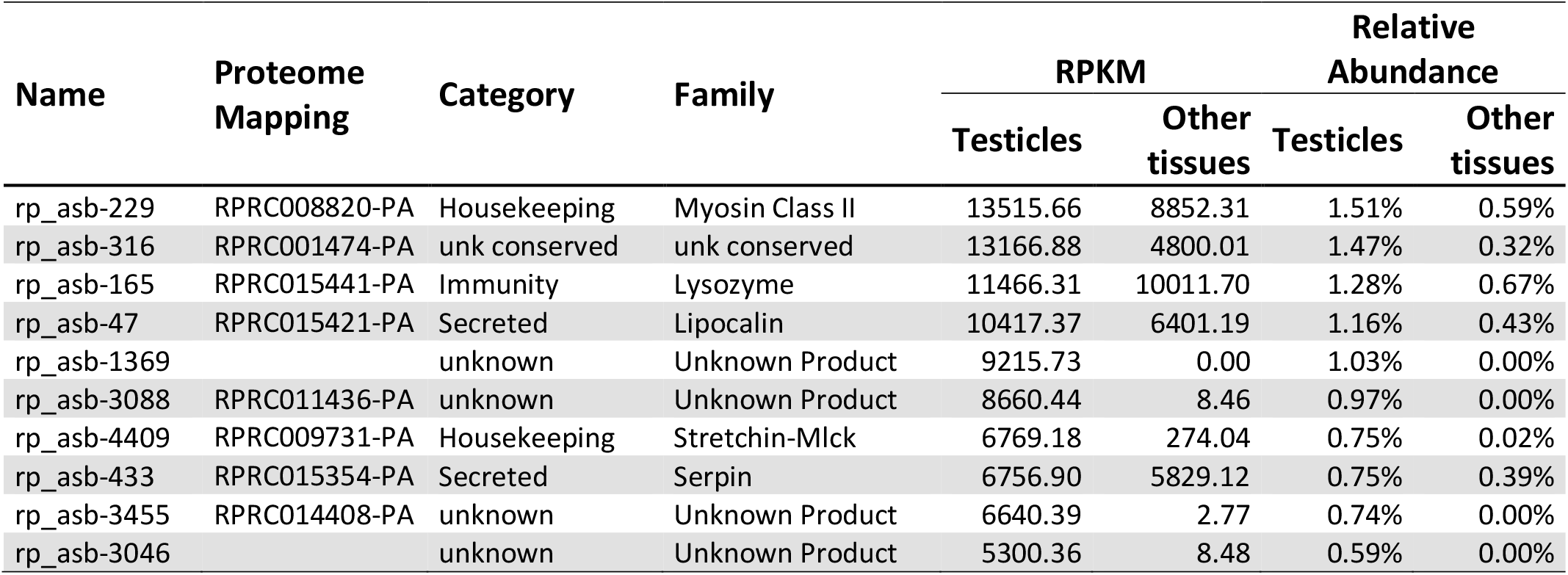
Top ten expressed genes in *Rhodnius prolixus* testicles.

| Name | Proteome Mapping | Category | Family | RPKM |  | Relative Abundance |  |
| --- | --- | --- | --- | --- | --- | --- | --- |
|  |  |  |  | Testicles | Other tissues | Testicles | Other tissues |
| rp_asb-229 | RPRC008820-PA | Housekeeping | Myosin Class II | 13515.66 | 8852.31 | 1.51% | 0.59% |
| rp_asb-316 | RPRC001474-PA | unk conserved | unk conserved | 13166.88 | 4800.01 | 1.47% | 0.32% |
| rp_asb-165 | RPRC015441-PA | Immunity | Lysozyme | 11466.31 | 10011.70 | 1.28% | 0.67% |
| rp_asb-47 | RPRC015421-PA | Secreted | Lipocalin | 10417.37 | 6401.19 | 1.16% | 0.43% |
| rp_asb-1369 |  | unknown | Unknown Product | 9215.73 | 0.00 | 1.03% | 0.00% |
| rp_asb-3088 | RPRC011436-PA | unknown | Unknown Product | 8660.44 | 8.46 | 0.97% | 0.00% |
| rp_asb-4409 | RPRC009731-PA | Housekeeping | Stretchin-Mlck | 6769.18 | 274.04 | 0.75% | 0.02% |
| rp_asb-433 | RPRC015354-PA | Secreted | Serpin | 6756.90 | 5829.12 | 0.75% | 0.39% |
| rp_asb-3455 | RPRC014408-PA | unknown | Unknown Product | 6640.39 | 2.77 | 0.74% | 0.00% |
| rp_asb-3046 |  | unknown | Unknown Product | 5300.36 | 8.48 | 0.59% | 0.00% |

**Figure 3.**
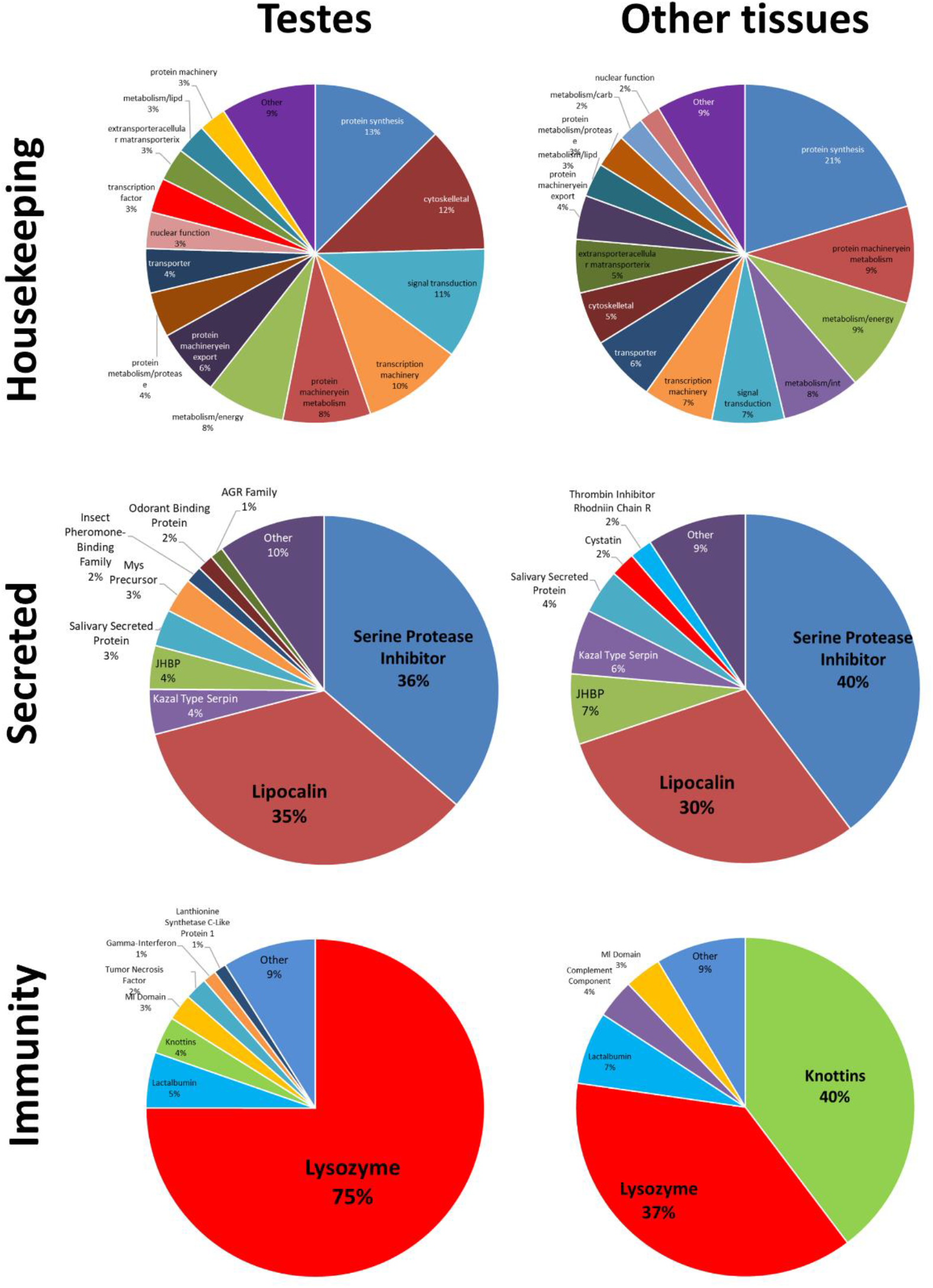
Relative abundance of transcripts by functional ontologies in three different major functional classes in testicles and other tissues of *Rhodnius prolixus*. Other tissues of *R. prolixus* are gut, ovaries, malpighian tubules, and fat body. Abundance is based in reads per kilobase million (RPKM).

#### Secreted

Our data suggests that the expression profile of secreted proteins in the testicles of *R. prolixus* is very similar to other tissues (Figure 3 and S3 Table), with lipocalins and serine protease inhibitors (serpins) as the most abundant families (corresponding to 71% of testicles and 70% of other tissues secreted transcripts). In hematophagous insects, lipocalins and serpins are usually associated to key salivary proteins that inhibit host hemostasis during blood ingestion. However, a few studies have demonstrated the role of lipocalins in sperm maturation and the role of serpins in spermatogenesis^14^. We found one lipocalin and one serpin amongst the most expressed genes in testicles (table 1), and an in-depth discussion on these families are detailed further in this manuscript.

#### Immunity

Immunity-related transcripts represents less than 2% of all transcripts expressed in testicles and other tissues (Figure 2). Nonetheless, our data suggests striking differences in the expression profile of testicles immunity related genes in comparison to other tissues (Figure 3). We observed that lysozymes correspond to 75% of all immunity related transcripts in testicles, while these proteins correspond to 37% of all immunity related transcripts in other tissues (Figure 3 and S4 Table). Lysozymes are enzymes with antimicrobial activity that recognize peptidoglycans.

Lysozymes have been found on seminal fluids of insects and mammals, but its role in this tissue remains to be elucidated. Discussion on *R. prolixus* testicles lysozymes are detailed further.

### Specific and differentially expressed genes in *Rhodnius prolixus* testicles

To have some insight on the regulation of gene expression of *R. prolixus* testicles, we performed a heatmap analysis using the mean z-scores for each gene family (Figure 4). Although the mean z-score can hide some overexpressed genes, this analysis avoids the complexity of evaluating the expression pattern of tens of thousands of genes, providing a simple and fast identification of gene families that may have important roles in tissue organization, spermatogenesis, sperm maturation, fertility, etc. Heatmaps were separated in the three major categories: secreted (Figure 4 panel A), housekeeping (Figure 4 panel B), and immunity (Figure 4 panels C1 and C2). As expected, we could not observe clear differences in the expression profile of housekeeping gene families (Figure 4 panel B), even in cytoskeletal transcripts, which have a relative abundance in testicles much higher than in other tissues. This result suggests that the abundance of cytoskeletal genes in testicles is a result of a few overexpressed genes. In fact, from the 634 cytoskeletal transcripts found in the transcriptome, only 429 are expressed in testicles, while 519 are expressed in other tissues (~20% more than testicles). On the other hand, we observed clear differences in the secreted and immunity categories. From the secreted gene families, our data suggests that the expression of mucins, anterior gradient (Agr), mys precursor, and serine protease inhibitors of the serpin and Kazal families are generally upregulated in testicles and down regulated in other tissues, while ubiquitin, triabin, apyrases, fringe-like proteins, salivary phospholipases, tryptophan-rich proteins, 47kda salivary proteins, inositol polyphosphatase, and platelet aggregation inhibitor proteins seem to be down regulated in testicles. As expected, many of the downregulated gene families in testicles are important salivary genes with a key role in inhibiting host hemostasis during blood feeding. In the immunity category (Figure 4 panel C1), we observed a general downregulation in lysozymes, tyrosine phosphatase, defensins, chemokines, and spatzle-3 gene families. It is interesting to note the down regulation of lysozymes in *R. prolixus* testicles, since transcripts of this family correspond to 75% of all immunity transcripts found in the testicles. The reason is that only two lysozymes out of nine are transcribed in testicles, and one of them is the third most abundant gene in the testicles’ transcriptome (table 1). Since the number of immunity related transcripts in the whole transcriptome is relatively low (exactly 100 genes), we decided to analyse the z-scores individually in this category (Figure 4 panel C2). From those, 12 genes are specifically expressed in testicles (Gamma-Interferon, Lanthionine Synthetase, Membrane Glycoprotein, 2 Phosphatases, Rhomboid, 4 Tumour Necrosis Factor genes, and 2 unclassified genes), and 34 genes are not expressed in testicles (mostly from the downregulated families). A more detailed discussion on testicles specific genes are detailed further in this manuscript.

**Figure 4.**
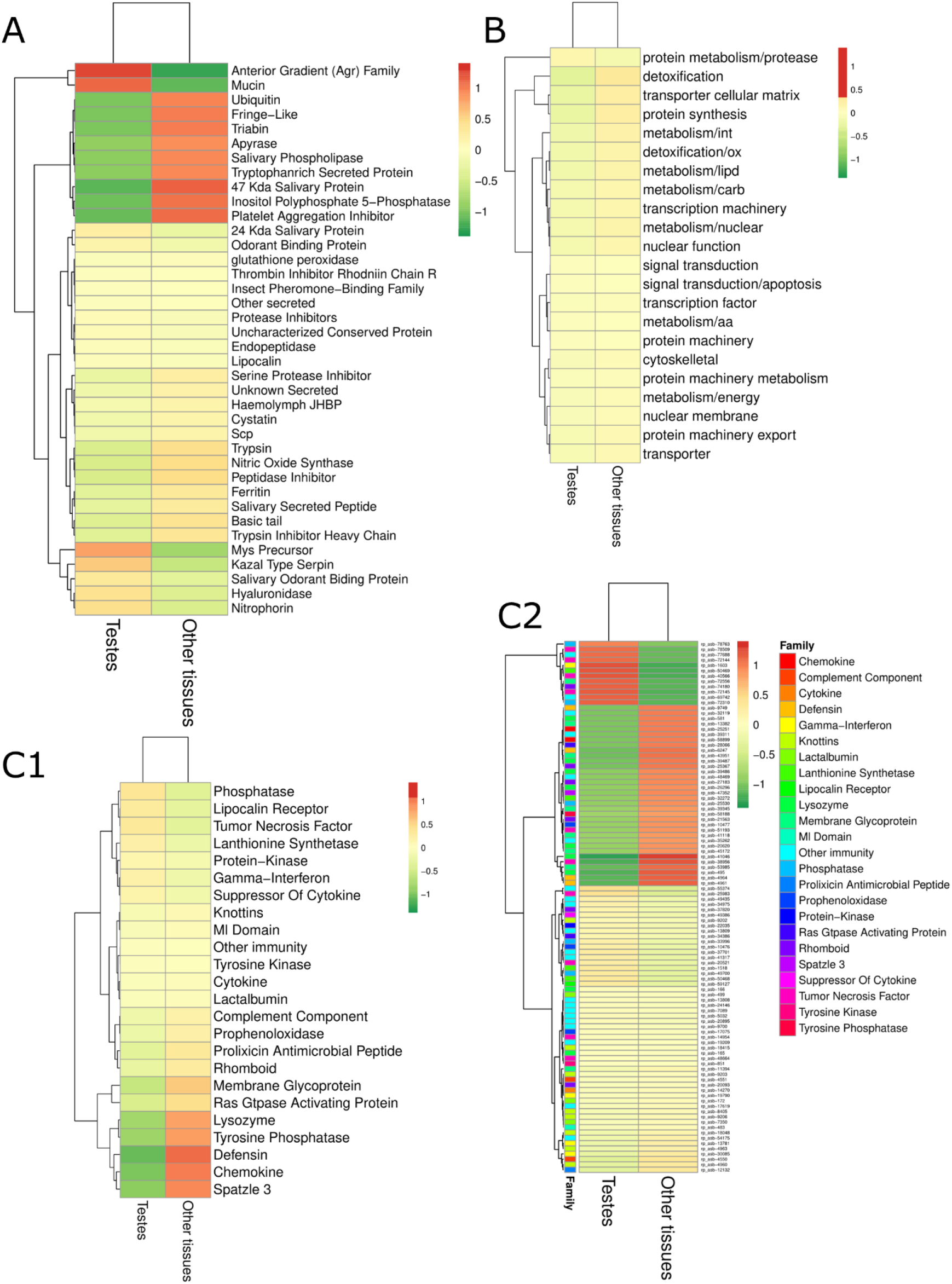
Heatmaps for each major functional category. Heatmaps show transcript abundancy (expression level) in each sample. **A** Secreted; **B** Housekeeping; **C1** Immunity; **C2** All immunity genes. In Bright Red are upregulated classes of genes (>2-fold standard deviation); Bright Green are downregulated classes of genes (< 2-fold standard deviation); Brownish green or red are classes with standard transcription. Classes were grouped by a pattern of transcription (eg. Genes that are upregulated in both tissues; genes that are differentially expressed; genes that are downregulated in both tissues; genes that have a standard regulation in both tissues; etc).

An interesting question regards the expression profile of uncharacterized genes (unknown and unknown conserved), since this group corresponds for more than seven thousand genes and for ~55% of transcript abundancy in testicles. From the 2,533 unknown conserved genes in the whole transcriptome, 431 have a testicles specific expression profile, while 856 are not expressed in testicles (S1 Excel Table). From those genes with expression in both testicles and other tissues, at least one seems to be upregulated in testicles (rp_asb-316/RPRC001474), with a RPKM value 3-fold higher in testicles than in other tissues (Table 1). From the 13,874 genes with unknown function and not conserved in other organisms, 3,563 contigs are specifically expressed in testicles, and 7,479 are not transcribed in testicles (S1 Excel Table). Four of those genes are amongst the top ten transcribed products in testicles (Table 1). From the 5,364 testicles-specific transcripts in this transcriptome, ~75% (3,994 transcripts) have unknown function and most of them could be novel *R. prolixus* proteins. Many proteomes and transcriptomes of the male reproductive organs describe the diversity of seminal fluid proteins produced by the male accessory glands, and we suspect that many of the testicles specific unknown proteins found here could belong to this class of proteins. Our reason to believe that is because such proteins are known to be species specific short peptides (many of them have a role in species compatibility) that evolve very fast. Indeed, the mean size of *R. prolixus* testicles specific peptides with unknown function is 100.68 amino acids, while the mean size of peptides in the whole transcriptome is 160.05 amino acids. Still, such claims should be answered in a further transcriptome aiming specifically to male accessory glands.

### Male fertility factors and other flagellar components expressed in testicles

Flagella are found in many prokaryotic and eukaryotic cells and are a typical component of sperm. In eukaryotes, the flagellum main structure is the axoneme, composed mainly by an intricate combination of microtubules (tubulins) and dyneins^15^. While microtubules are responsible for support, dyneins act as motors, giving movement to the flagellum^16^. There are several isoforms of axonemal dynein heavy chains (α, β, γ, 1-α, 1-β, etc.) that associate to form the inner and outer arms of the axonemes. In *D. melanogaster*, the Y chromosome harbours three dynein heavy chains known as *kl-2*, *kl-3,* and *kl-5*, and knockout of any of these genes causes male sterility due to lack of flagellum movement^17–19^. We found 45 tubulins and 30 dyneins in the *R. prolixus* transcriptome. From those, 19 tubulins and 15 dyneins presented testicles specific expression, and only six tubulins and three dyneins were not expressed in testicles (S1 Excel Table). Tubulins are 3-fold more expressed in testicles than in other tissues, while dyneins are 20-fold more expressed in testicles. We also found that all transcripts mapped to the insect genomic scaffolds, but only 18 mapped to annotated proteins. Evolutionary analysis of the dynein heavy chains (Figure 5) suggests that *R. prolixus* have two orthologs of *D. melanogaster kl-2* (which belongs to the dynein heavy chain 1-β family), and one of them has a testicles specific expression pattern. However, none of them are amongst the most expressed dyneins in the transcriptome. The two most expressed dyneins in testicles are rp-asb-15029 (unmapped to proteome) and rp_asb-59266/RPRC003396 (RPKM 397.39 and 177.55 respectively). Rp-asb-59266 is not specific to testicles (it is also expressed in the anterior and posterior midgut) but is expressed ~66-fold in testicles than in the other tissues. On the other hand, transcript rp_asb-59266/RPRC003396 is specific to testicles and the structural and evolutionary analysis suggests it is a dynein light chain with two isoforms. Interestingly, we did not find any orthologs for *kl-3* and *kl-5*, which are well conserved genes in eukaryotes, arising a question if these transcripts were not detected in our study or if such genes were lost on the *R. prolixus* genome (we searched for sequences in the genomic and transcriptome raw reads without success). Also, six of the testicles specific dyneins showed no evolutionary similarities with other dyneins with known function. Hence, further studies on the role of dyneins in the fertility of kissing bugs are needed to better understand the role of these proteins in male fertility.

**Figure 5.**
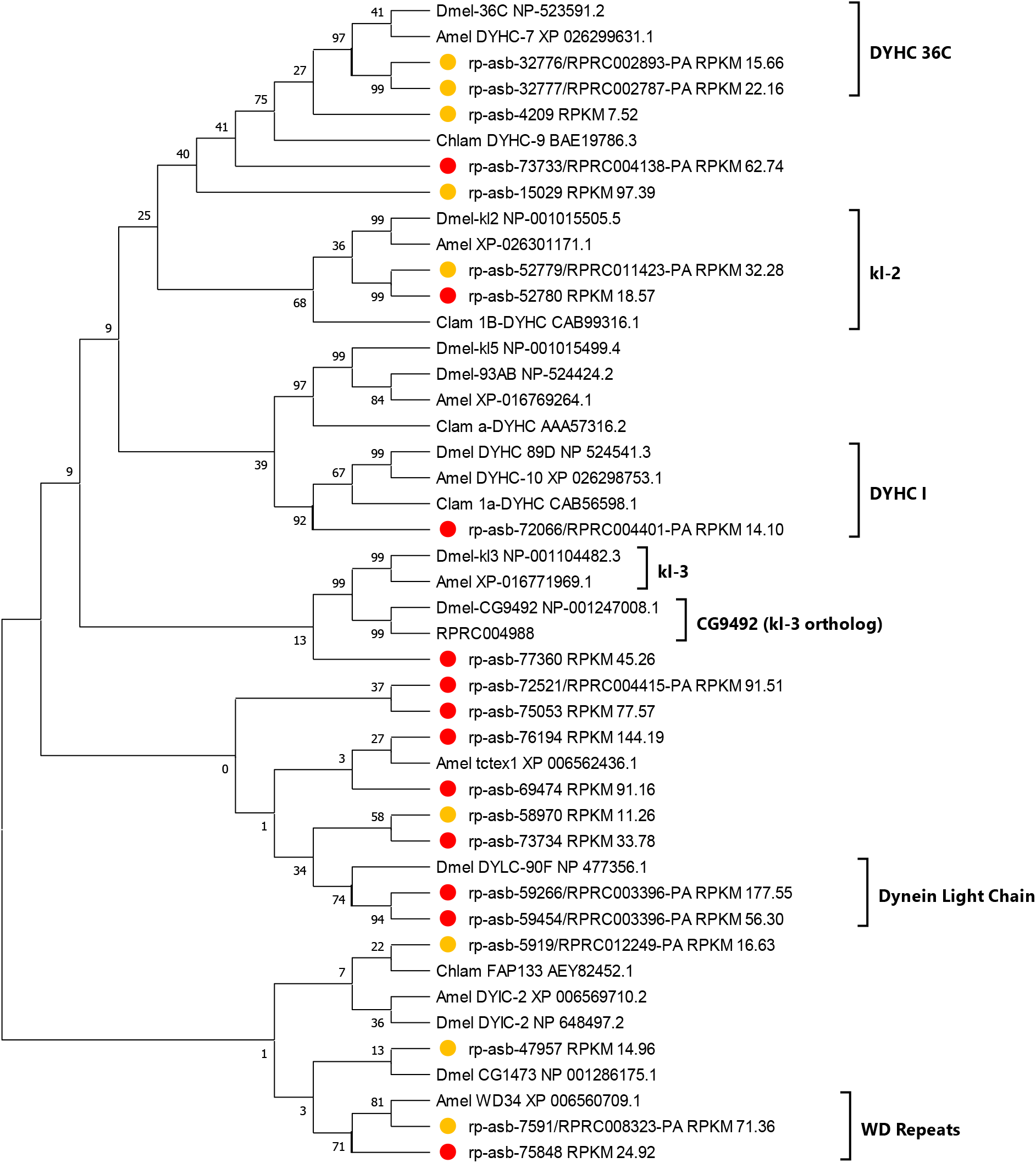
Evolutionary relationships of *R. prolixus* dynein proteins. The evolutionary history of the *R. prolixus* dyneins was inferred using the Neighbor-Joining method. Red dots show transcripts with testicles specific expression pattern, and yellow dots show non-specifc expression pattern. RPKM values of each transcript are shown in each branch, and arrows indicate transcript with the highest RPKM values. Accession numbers of non-*Rhodnius* proteins are shown in each branch after species abbreviation. Species abbreviations are: Dmel = *Drosophila melanogaster*; Amel=*Apis mellifera*; Chlam=*Chlamydomonas* sp. Evolutionary analyses were conducted in MEGA.

**Figure 6.**
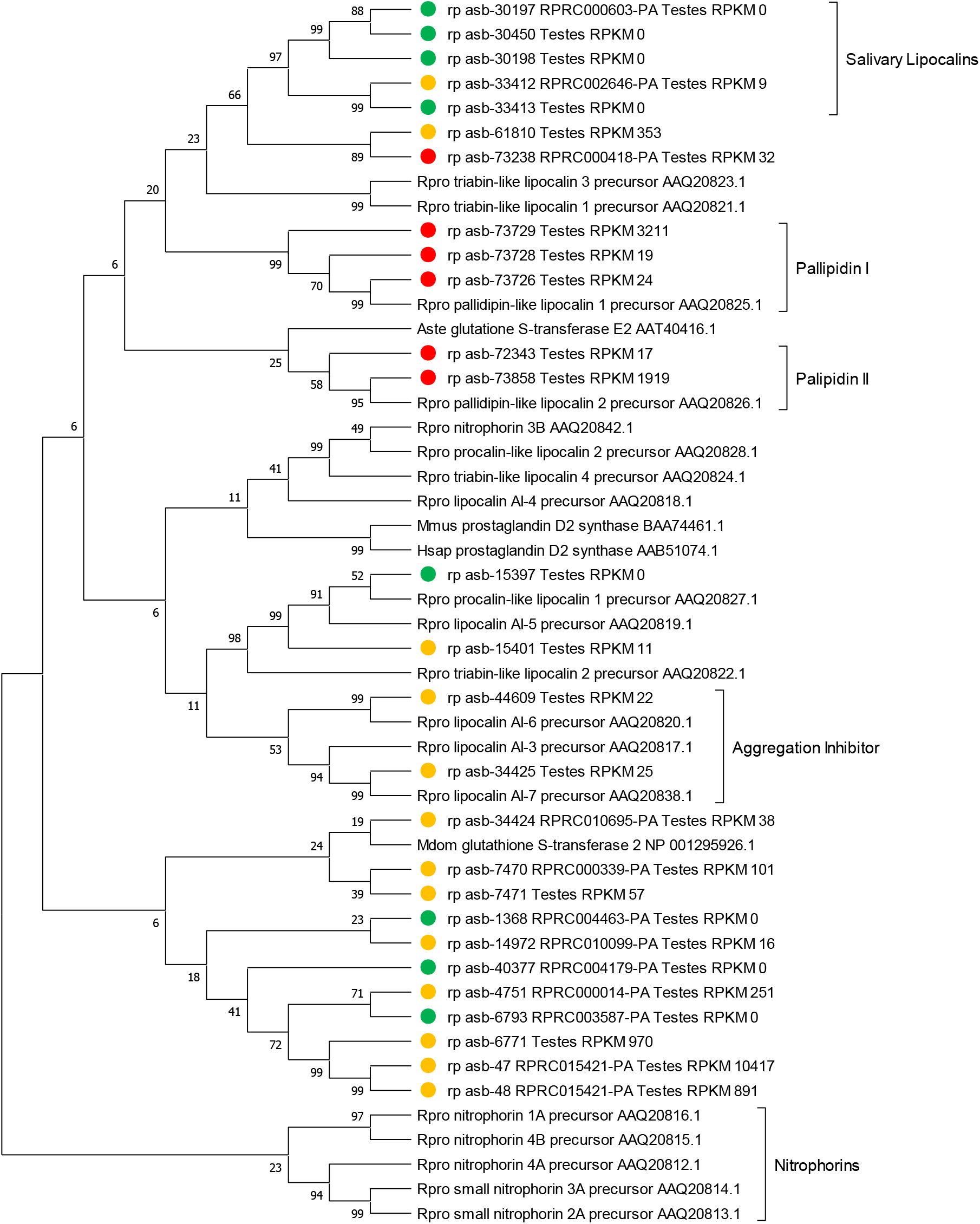
Evolutionary relationships of *R. prolixus* lipocalins. The evolutionary history of the *R. prolixus* lipocalins was inferred using the Neighbor-Joining method. Yellow dots show transcripts with testicles non-specific expression pattern. Green dots show transcripts not expressed in testicles. RPKM values of each transcript are shown in each branch and arrows indicate transcripts with the highest RPKM values. Accession numbers of non-*Rhodnius* proteins are shown in each branch after species abbreviation. Evolutionary analyses were conducted in MEGA.

### Spermatogenesis related genes

Spermatogenesis is an interesting phenomenon found in eukaryotes and is a predominant part of sexual reproduction. The process begins with mitotic divisions of spermatogonia stem cells to create spermatocyte cells^20^. Spermatocytes suffer meiotic division to form haploid cells that will maturate (during spermiogenesis) into spermatozoa. Despite its importance, mechanisms underlying spermatogenesis (specially the meiotic process) remain largely obscure. There are two characteristics in kissing bugs spermatogenesis that make this process even more intriguing. First is the fact that kissing bugs chromosomes are holokinetic (they do not have a centromere), and the processes of segregation of holokinetic chromosomes during cell division remain unclear^21,22^. The second characteristic is that in kissing bugs the meiosis of sex chromosomes occurs in an inverse order, in which sister chromatids segregate in the first stage of meiosis and homologs segregate in the second stage^23^. We identified three gene families that are well known for their role in spermatogenesis: myosin, actin, and tumour necrosis factors (TNF).

Myosin is a superfamily of actin-dependent molecular motor proteins known for their roles in muscle contraction in eukaryotes. These proteins also have a key role in spermatogenesis, such as acrosome biogenesis, spindle assembly and positioning, karyokinesis, and spermatid individualization^24^. We found 114 expressed transcripts from the myosin family, from which 19 presented a testicles specific expression pattern (S1 Excel Table). The most abundant transcript in testicles is the myosin rp-asb-229/RPRC008820 (Table 1), and it has non-specific expression pattern, being also very abundant in gut tissues^25^ (S1 Excel Table). However, the second and third most expressed myosin in testicles (rp-asb-70879/RPRC010999 and rp-asb-71794/RPRC000888 respectively) are testicles-specific, and it would be very interesting to further evaluate their role in spermatogenesis using silencing (RNAi) techniques. On the other hand, only one actin (rp-asb-71413) from the 44 identified actins is specifically expressed in testicles. Actins are critical for correct nuclear positioning, germinal vesicle breakdown, spindle migration, spindle rotation, and chromosome segregation in spermatogenesis^24^. Hence, the question on the role of this testicles specific actin remains open.

The specific role of TNFs in spermatogenesis of insects is still unknown. TNFs are cytokines involved in the regulation of immune cells. It can induce fever, apoptotic cell death, inflammation, inhibition of tumorigenesis, etc. Many studies have demonstrated that disruption of TNF expression in testicles can affect spermatogenesis, imputing these molecules as essential for maintaining germ cell homeostasis and functional spermatogenesis in testicles^26,27^. In mammals, testicles specific TNF-alpha has an anti-hormonal role and play interactions between Sertoli and germ cells (Sertoli cells are thought to play a central role in male-specific cell interactions, including those that occur during spermatogenesis)^28^. In Drosophila, the testicles contains two types of stem cells, germline stem cells (GSCs) and cyst stem cells (CySCs)^29^, and it remains to be seen whether *R. prolixus* has the same population of germline stem cells. From the ten *R. prolixus* TNFs identified in the transcriptome, we found that rp_asb-20521 is differentially expressed in testicles, and four (rp_asb-40566, rp_asb-72145, rp_asb-72144/RPRC007130, rp_asb-78509/RPRC003901) that have a testicles specific expression pattern (Figure 4, C2). Such genes could be targets to understand the role of TNF in insects and its interaction, if any, with GSCs and CySCs.

### Secreted lipocalins

The lipocalins are a family of proteins which transport small hydrophobic molecules such as steroids, bilins, retinoids, and lipids^30^. They share limited regions of sequence homology and a common tertiary structure architecture^31^. Lipocalins have been associated with many biological processes, among them immune response, pheromone transport, biological prostaglandin synthesis, retinoid binding, and cancer cell interactions^30^. Ticks and kissing bugs evolved salivary lipocalins that act as efficient scavengers of biogenic amines^32–34^ and eicosanoids^35^.

Lipocalins represented 35% of all secreted transcripts in testicles, and from the 19 lipocalins identified in *R. prolixus* transcriptome, we found that eight (42%) are specifically expressed in testicles, while only five are not expressed in testicles. The lipocalin rp_asb-47 is the fourth most abundant transcript in testicles, and it is nearly 2-fold more abundant in testicles than in other tissues. Interestingly, none of the testicles specific lipocalins are highly expressed (RPKM values bellow 50). Phylogenetic analysis comparing the transcriptome lipocalins with lipocalins from GenBank suggests that a few testicles specific lipocalins could have general roles in platelet aggregation inhibition: pallipidin (rp_asb-73726, rp_asb-73728, rp_asb-73729), triabin/procalin (rp_asb-15401), and platelet aggregation inhibitors (rp_asb-34425). One could wonder why inhibitors of platelet aggregation would be specifically expressed in *R. prolixus* testicles. First, in hematophagous insects, lipocalins have been mostly studied in the light of the blood feeding process, which creates a bias in our knowledge. Secondly, it is important to note that salivary glands (SG) were not included in this transcriptome (the reason was that SG had already been extensively studied before). However, if these lipocalins are produced in the accessory glands and transferred to the female spermatheca, they could modulate the female smooth muscle contractibility by scavenging or transferring biogenic amines and/or eicosanoids. A separated transcriptome of the accessory glands with a testes transcriptome should clarify these hypotheses. Lipocalins have been detected in the semen of other organisms. However, their role in male fertility is incipient. The best known testicles specific lipocalins are the prostaglandin D-synthase lipocalins of mammals^36^, where it is hypothesised that lipocalins have a role as carriers of hormones and retinoids to the developing germ cells in the seminiferous tubules and the maturing spermatozoa^14^. Alternatively, it is possible that some of the testicles specific lipocalins that are also expressed in SG have no analogous role. Interestingly, all three lipocalins that showed higher expression in testicles (rp_asb-47/RPRC015421, rp_asb-61810, rp_asb-7470/RPRC000339) do not correlate with lipocalins with known function. Although these genes are not exclusively expressed in testicles, it is possible that they might have a central role in sperm maturation with a role not yet known to insect lipocalins (such as the transfer of prostaglandins and eicosanoids during mating^7^), turning them into important targets for further functional studies.

### Secreted serine protease inhibitors

Serpins transcripts constitute ~36% of all secreted proteins in testicles. Serpins are a superfamily of proteins with similar structures that were first identified for their protease inhibition activity and are found in all kingdoms of life. In the *R. prolixus* transcriptome, we found 36 serpins, but none of them presented a testicles specific expression pattern. Twenty-three of them seem to be highly, and equally, expressed in testicles and gut (mean RPKM of 611.4 in testicles and 498.7 in gut). Serpins have been detected in turkey testicles, epididymis, ductus deferens, spermatozoa surface, and in seminal plasma using electrophoretic methods^37^. From the 36 serpins found, transcript rp_asb-21159 caught our attention due to its differential expression in relation to other tissues (~25-fold higher). However, when we looked for known domains in rp_asb-21159/RPRC014952 sequence, we found that it contains a well-known 7tm-odorant binding domain, suggesting a miss annotation in the transcriptome. Hence, although serpins are highly abundant in testicles, we could not identify specific contigs or patterns that differentiate from other tissues, and the question on the role of serpins in male reproduction remains open ended. In addition to serpins, Kazal-family serine protease inhibitors are found in seminal plasma, known as acrosin inhibitors. A study showed that serine protease inhibitor Kazal-type 2 (SPINK) is required for maintaining normal spermatogenesis and potentially regulates serine protease-mediated apoptosis in male germ cells^38^.

### Lysozymes

Lysozymes are antimicrobial enzymes produced by animals that form part of the innate immune system. These proteins have a specific role in the hydrolysis of N-acetyl-D-glucosamine residues in peptidoglycan, causing lysis of bacteria. Lysozymes are abundant in secretions, including tears, saliva, human milk, and mucus. *R. prolixus* transcriptome suggests that lysozymes may have an important role in testicles, representing 75% of all immunity related transcripts. Recently, testicles specific lysozyme-like genes (Lyzl) belonging to the c-type lysozyme family have been described in sperm proteome of humans and mouse. However, their role in male fertility and mammal reproduction remains unclear. In the bed bug *Cimex lectularius*, the antimicrobial activity of the seminal fluid is attributed to Lyzl, and it was hypothesized that these genes could have a role in helping to protect the female reproductive tract from bacteria introduced during copula. From the nine lysozyme transcripts identified in *R. prolixus* transcriptome, only two of them, rp-asb-165/RPRC15441-PA and rp-asb-166/RPRC15441-PA are expressed in testicles (both are expressed in other tissues, mainly gut). Interestingly, both lysozymes map to the same protein RPRC15441, initially suggesting they are splice variants from the same gene. However, in a closer inspection, lysozyme rp-asb-165 is much longer than rp-asb-166 due to an extension of nearly 600 nucleotides in the 3’ end. Using blast searches to understand both lysozymes structures, we found that 3’ extension could be the result of a chimera of the transcriptome assembly, since its 300 final nucleotides shows homology to other scaffold with 96% nucleotide identities (Figure 7a). Additionally, there is a region of ~300 nucleotides in the middle of the transcript that do not present homology to any assembled scaffold. Hence, it is more likely that rp-asb-165 is a result of a miss-assembly. Still, evolutionary analysis (Figure 7b) also suggests testicles lysozymes are orthologous to the lysozyme duplications *Lys* of *Drosophila melanogaster* (Figure 7b). Interestingly, we found that most of the non-testicles lysozymes are in members of the peptidoglycan recognition proteins LC family (PGRP-LC). These transmembrane proteins act as signal transducing receptors, initiating the immune cascade by recognition of bacterial peptidoglycans followed by activation of the innate immunity *Imd* pathway.

**Figure 7.**
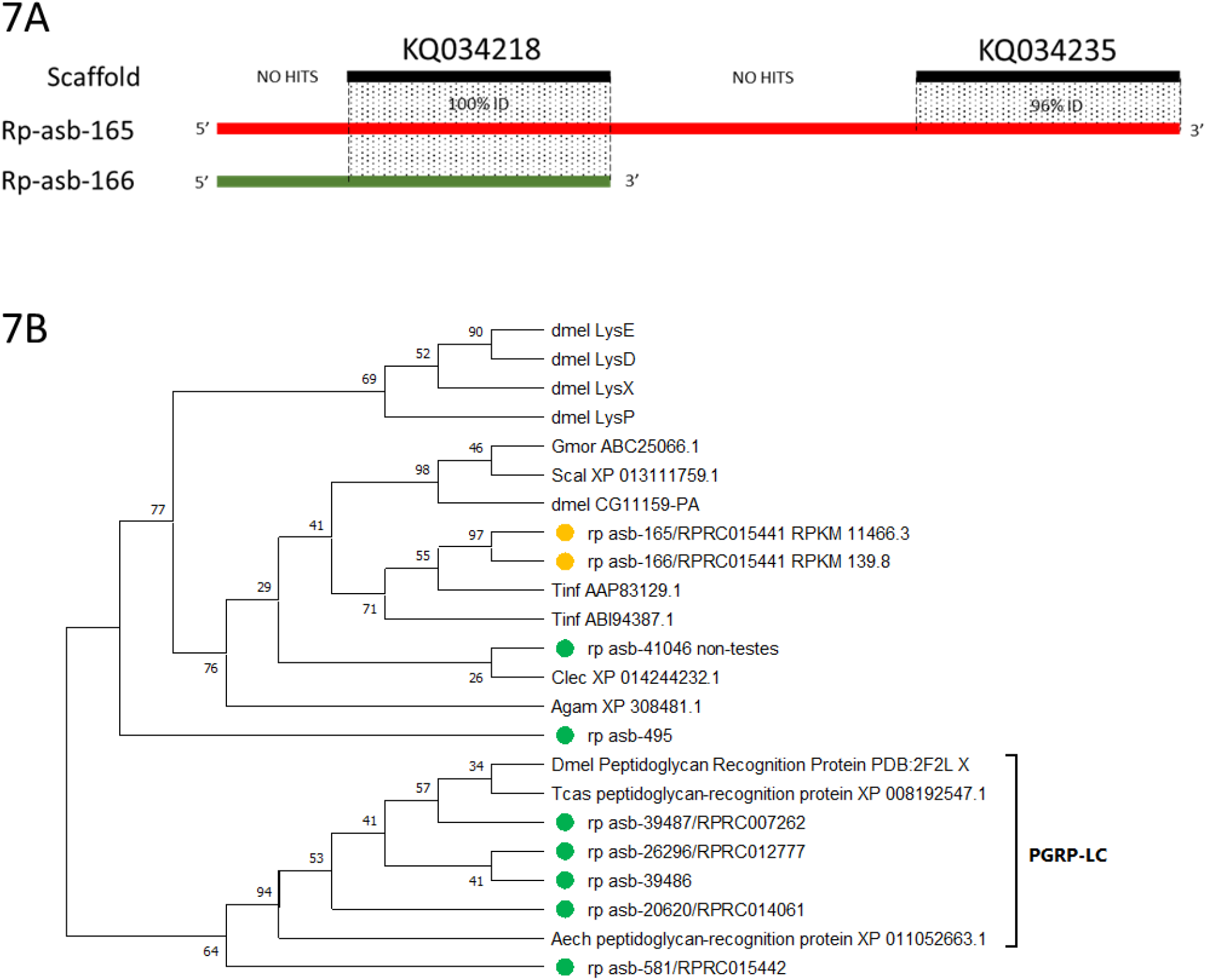
Evolutionary relationships of *R. prolixus* lysozymes. **7A** shows the structural analysis of testicles expressed lysozymes rp-asb-165 and rp-asb-166 ascertained by blast alignments to the genomic scaffolds. The shadowed area shows the homologous regions of each transcript to scaffolds (black lines), including the nucleotide identity between scaffold and transcripts. **7B** shows the evolutionary history of the *R. prolixus* lysozymes which was inferred using the Neighbor-Joining method. Yellow dots show non testicles-specific expression pattern, and green dots show transcripts not expressed in testicles. RPKM values of each transcript are shown in each branch. Accession numbers of non-*Rhodnius* proteins are shown in each branch after species abbreviation. Species abbreviations are: Dmel = *Drosophila melanogaster*; Scal= *Stomoxys calcitrans*; Tinf=*Triatoma infestans*; Agam=*Anopheles gambiae*; Clec=*Cimex lectularius*; Tcas = *Tribolium castaneum*; Aech = *Acromyrmex echinatior.* Evolutionary analyses were conducted in MEGA.

### Mapping the transcriptome to *Rhodnius prolixus* genome and proteome

When the sequencing and assembling of *R. prolixus* transcriptome finished, in 2014^25^, the insect genome was still under analysis and was published only in 2015^13^. Hence, the transcriptome and the first manuscript about its data were not linked to the insect’s genome. Also, although the genome annotation used this transcriptome as reference for protein prediction, the published genome was never linked to the transcriptome. This is unfortunate because the transcriptome contains valuable data on differential tissue expression, allowing much deeper studies and insights on the *R. prolixus* physiology. Therefore, we decided to join both, transcriptome and genome, to create a comprehensive information on the genomic functionality of the kissing bug *R. prolixus* (S1 Excel Table). More than 85% of the transcripts mapped to the insect genome (22,052 of 25,673) with identity above 98% (S1 Table). The distribution of unmapped transcripts was mostly even among categories (from 4% to 20% of unmapped transcripts), and the category with most unmapped transcripts was transposable elements, with ~25% of unmapped transcripts (which is expected since most TEs are inserted in heterochromatic/repetitive regions, which is not assembled in final genomes^39^, or their annotation are purposefully avoided as “repeats”). We observed that many transcripts mapped to the same genomic coordinates, suggesting the presence of genes with multiple splice variants. In a manual observation, we counted at least 2,000 possible splice variants (it is not the scope of this manuscript to endeavour on splicing variation on *R. prolixus* and we will not discuss this issue in depth. Still, as far as we know, this is the first observation of splicing variants in *R. prolixus* genome). On the other hand, only 36% of the transcripts mapped with more than 98% to the annotated genome coding sequences of *R. prolixus* (9,311 out of 25,673), which is very surprising. In fact, when we look at transcripts that aligned to different proteins, only 7,126 different proteins mapped to the transcriptome (2,185 transcripts aligned to same proteins, suggesting splice variants). It is important to note that the genomic strain of *R. prolixus* is from a certified colony at CDC Atlanta (USA), while the strain used for the transcriptome is from a colony from the Federal University of Rio de Janeiro (Brazil). To test if strain divergence was responsible for the low mapping success, we observed the effect of lower identities on blast hits (S5 Table). Still, even with a cut-off of 50% on amino-acid identities, only 8,905 transcripts mapped to different proteins (which is ~60% of the *R. prolixus* proteome). For a perspective, in *Drosophila* species, amino-acid identities between genes from species separated for 1.5 million years (*D. melanogaster* and *D. simulans*) is above the 70% level^40^. It is important to note that in our research we found many miss-assemblies in the transcriptome (which could be a result of the sequencing technology of choice), which could explain why so many transcripts do not hit to any proteins. Nonetheless, we found that 3,297 transcripts mapped to scaffolds with 100% nucleotide identity and failed to align to any proteins with more than 98% amino-acid identity (S6 Table). Exploring such regions, we found that no proteins were annotated in these genomic regions. Looking at the mapping success rate by functional roles (Supplementary Tables 1-6), while housekeeping, immunity, secreted, and unknown conserved transcripts mapped to the proteome at a reasonable rate (between 54% and 70%), more than 85% of the transcripts without known homologs (unknown category) did not map to *R. prolixus* annotated proteome. As discussed before, most of the transcripts from the unknown category code for very short peptides (less than 100 amino acids), which could explain why the annotation process failed to identify these putative genes.

In summary, our mapping effort strongly suggest that the annotated *R. prolixus* proteome can be under-estimated in at least 3,000 proteins. We know that the *R. prolixus* genome is very fragmented (which could also explain the loss of nearly 3,000 proteins), and given the importance of this insect to public health and research in insect physiology, it is imperative that a re-sequencing of the genome (using the new ultra-long read sequencers) and a re-annotation of the proteome must be performed.

### Final insights on the *Rhodnius prolixus* testicles transcriptome

Although *R. prolixus* is a model for studies on insect physiology, the knowledge on the genetics of this species has lagged behind. In such cases, transcriptome studies have the power of creating a catalogue of candidate genes to be further investigated to better understand insect genetics. The results found on *R prolixus* transcriptome is a reflection on our lack of knowledge in triatomine genetics, in which the function of more than 60% of the annotated transcripts remains unknown. In triatomines, most studies aim to better understand the hematophagy process, and although the *R. prolixus* transcriptome was generated from a multitude of tissues, the paper describing *R. prolixus* transcripts focused on the physiology of gut. Therefore, in our study we evaluated the transcripts from *R. prolixus* testicles, and this is the first study focused on *R. prolixus* male reproduction in the genomic era. As expected, we do not have a clue on the function of ~50% of the testicles’s transcripts; and five of the top ten most expressed transcripts in testicles belong to this category. However, even the role for these transcripts containing well described domains in reproduction remains unclear. This is also the case for genes well studied in *R. prolixus* physiology, such as lipocalins, serpins, and lysozymes. These three families of genes have important roles in the blood-feeding process, and surprisingly, these were the most abundant families of secreted and immunity related transcripts. Interestingly, mosquitoes have an expanded salivary family, named the D7 family^41–43^, containing one or two odorant-binding motifs that inhibit hemostasis by binding biogenic amines and leukotrienes, a function similar to the salivary lipocalins of *Rhodnius.* A study of the accessory glands of the mosquito, *Aedes aegypti,* identified the presence of one typical highly expressed salivary D7 protein^44^. It is thus possible to speculate that many genes co-opted by evolution for expression by the salivary glands of hematophagous arthropods may have their origins as male testicles, or accessory glands-expressed genes. It is possible that such genes have a role in testicles very similar (regarding their molecular ligands) to their counterparts expressed in the salivary glands or gut, but it is also very possible that lipocalins, serpins, and lysozymes specifically expressed in testicles have distinct roles that we still do not understand. On the other hand, the testicles transcriptome generated a catalogue of transcripts with well described roles on spermatogenesis and insect fertility. We identified myosins, dyneins, actins, tumour necrosis factors and other genes that could disrupt spermatogenesis and cause male sterility. Such genes should be targets on functional studies to better understand triatomine male biology, a field of study that is mostly neglected by the scientific community.

## Acknowledgments

We would like to thank Brian Brown, NIH Library Editing Service, for reviewing the manuscript. This research is part of the Brazilian research program Institutos Nacionais de Ciência e Tecnologia – Entomologia Molecular.

## Author contributions

We declare that this work was done by the authors named in this article and all liabilities pertaining to claims relating to the content of this article will be borne by the authors. All authors wrote, reviewed and approved the manuscript including figures and tables. L.B.K., P.L.O. and J.M.R. conceived the study. L.B.K., P.L.O., D.M., M.M. and M.H.S. prepared samples for sequencing. JMR and RDM. assembled and annotated the transcriptome. J.C., D.V.S., R.N.A., M.R.V.S., N.F.G., M.H.P., G.D.P. and L.B.K contributed to testicles transcriptome analysis. J.C. conducted abundance, heat-map and evolutionary analyses. J.C. and LBK edited the manuscript.

## Supporting Information

S1 Excel Table

S1 Table: Summary of transcripts by major functional classes.

S2 Table: Summary of housekeeping gene transcripts by functional protein families.

S3 Table: Summary of immunity gene transcripts by functional protein families.

S4 Table: Summary of secreted gene transcripts by functional protein families. S5 Table: Transcriptome mapping to proteome summary

S6 Table: number of transcripts, by functional groups, that mapped to genome with 100% nucleotide identity but failed to map to proteome with >98% amino acid identity

## Data availability

All sequences and annotation tables are freely available in the GenBank or as the Supplementary dataset.

