## Supplementary material for "Analysis of the testicle’s transcriptome of the Chagas disease vector *Rhodnius prolixus*": S1 Table

### S1 Table: Summary of transcripts by major functional classes.

| Major functional classes | # of transcripts | Expressed in Testes | Expressed in other tissues | Mapped to genome | Mapped to proteome | RPKM Testes | RPKM other tissues |
|---|---|---|---|---|---|---|---|
| Housekeeping | 8513 | 6012 | 7233 | 7936 | 5587 | 341622,50 | 380387,85 |
| Immunity | 100 | 66 | 88 | 96 | 70 | 15405,34 | 27765,52 |
| Secreted | 257 | 182 | 214 | 207 | 139 | 37736,74 | 28775,75 |
| Transposable element | 374 | 112 | 338 | 283 | 61 | 3225,67 | 10036,20 |
| unknown conserved | 2533 | 1677 | 2102 | 2366 | 1494 | 96027,46 | 92256,03 |
| unknown | 13874 | 6395 | 10311 | 11143 | 1956 | 402503,76 | 948291,65 |
| viral | 22 | 10 | 22 | 21 | 4 | 329,16 | 951,02 |
| **Total** | **25673** | **14454** | **20308** | **22052** | **9311** | **896850,64** | **1488464,01** |
