## Supplementary material for "Analysis of the testicle’s transcriptome of the Chagas disease vector *Rhodnius prolixus*": S2 Table

### S2 Table: Summary of housekeeping gene transcripts by functional protein families.

| Housekeeping protein functional classes | # of transcripts | Expressed in Testes | Expressed in other tissues | Mapped to genome | Mapped to proteome | RPKM Testes | RPKM other tissues |
|---|---|---|---|---|---|---|---|
| Cytoskelletal | 634 | 429 | 516 | 602 | 390 | 41104,54 | 19438,51 |
| Detoxification | 94 | 58 | 89 | 88 | 64 | 1986,16 | 2734,78 |
| Detoxification/ox | 109 | 73 | 91 | 95 | 61 | 2290,15 | 5786,09 |
| Extransporteracellular matransporterix | 309 | 175 | 267 | 291 | 191 | 10730,14 | 19384,02 |
| Metabolism/aa | 167 | 128 | 140 | 157 | 135 | 5348,39 | 5217,13 |
| Metabolism/carb | 265 | 191 | 232 | 254 | 183 | 7299,81 | 9332,02 |
| Metabolism/energy | 353 | 275 | 309 | 294 | 232 | 25623,96 | 34400,98 |
| Metabolism/int | 127 | 83 | 110 | 117 | 100 | 3929,25 | 28709,28 |
| Metabolism/lipd | 364 | 248 | 313 | 341 | 243 | 9968,58 | 12258,58 |
| Metabolism/nuclear | 176 | 124 | 157 | 161 | 116 | 3765,86 | 4355,73 |
| Nuclear function | 436 | 288 | 379 | 415 | 297 | 11885,98 | 7702,93 |
| Nuclear membrane | 30 | 26 | 30 | 29 | 21 | 3743,90 | 2502,73 |
| Protein machinery | 255 | 193 | 215 | 241 | 184 | 8875,06 | 5237,79 |
| Protein machinery export | 442 | 334 | 397 | 415 | 310 | 21830,47 | 16084,16 |
| Protein machinery metabolism | 503 | 391 | 440 | 455 | 349 | 28606,83 | 35319,89 |
| Protein metabolism/protease | 292 | 223 | 205 | 270 | 192 | 14679,83 | 12240,16 |
| Protein synthesis | 653 | 470 | 628 | 581 | 411 | 42902,71 | 78087,63 |
| Signal transduction | 1297 | 910 | 1010 | 1235 | 791 | 36107,32 | 26358,66 |
| Signal transduction/apoptosis | 87 | 65 | 71 | 83 | 60 | 2724,62 | 1672,40 |
| Transcription factor | 259 | 179 | 206 | 243 | 151 | 11276,32 | 5099,66 |
| Transcription machinery | 1054 | 746 | 942 | 1006 | 683 | 32766,21 | 25648,28 |
| Transporter | 607 | 403 | 486 | 563 | 423 | 14176,42 | 22816,43 |
| **Housekeeping total** | **8513** | **6012** | **7233** | **7936** | **5587** | **341622,50** | **380387,85** |
