## Supplementary material for "Analysis of the testicle’s transcriptome of the Chagas disease vector *Rhodnius prolixus*": S3 Table

### S3 Table: Summary of immunity gene transcripts by functional protein families.

| Immunity protein functional classes | # of transcripts | Expressed in Testes | Expressed in other tissues | Mapped to genome | Mapped to proteome | RPKM Testes | RPKM other tissues |
|---|---|---|---|---|---|---|---|
| Chemokine | 2 | 0 | 2 | 1 | 2 | 0,00 | 15,29 |
| Complement Component | 2 | 2 | 2 | 2 | 2 | 56,79 | 1031,60 |
| Cytokine | 1 | 1 | 1 | 1 | 1 | 65,14 | 124,23 |
| Defensin | 4 | 0 | 4 | 4 | 2 | 0,00 | 89,34 |
| Gamma-Interferon | 4 | 4 | 3 | 4 | 4 | 198,83 | 410,51 |
| Knottins | 8 | 8 | 8 | 7 | 7 | 553,04 | 11027,09 |
| Lactalbumin | 3 | 3 | 3 | 2 | 2 | 821,53 | 1921,09 |
| Lanthionine Synthetase | 4 | 3 | 3 | 4 | 3 | 182,64 | 7,48 |
| Lipocalin Receptor | 1 | 1 | 1 | 1 | 1 | 16,78 | 0,55 |
| Lysozyme | 9 | 2 | 9 | 9 | 6 | 11606,11 | 10424,61 |
| Membrane Glycoprotein | 8 | 2 | 7 | 8 | 6 | 64,63 | 164,55 |
| Ml Domain | 1 | 1 | 1 | 1 | 1 | 393,58 | 976,70 |
| Other immunity | 21 | 17 | 19 | 21 | 13 | 680,14 | 408,68 |
| Phosphatase | 5 | 4 | 3 | 5 | 5 | 124,35 | 7,06 |
| Prolixicin Antimicrobial Peptide | 1 | 1 | 1 | 0 | 0 | 24,67 | 882,22 |
| Prophenoloxidase | 3 | 2 | 3 | 3 | 3 | 27,33 | 13,03 |
| Protein-Kinase | 1 | 1 | 1 | 1 | 1 | 33,28 | 8,91 |
| Ras Gtpase Activating Protein | 2 | 1 | 2 | 2 | 0 | 69,25 | 22,63 |
| Rhomboid | 6 | 3 | 5 | 6 | 5 | 48,87 | 28,14 |
| Spatzle 3 | 1 | 0 | 1 | 1 | 1 | 0,00 | 3,60 |
| Suppressor Of Cytokine | 1 | 1 | 1 | 1 | 0 | 7,95 | 1,31 |
| Tumor Necrosis Factor | 10 | 8 | 6 | 10 | 4 | 333,15 | 93,53 |
| Tyrosine Kinase | 1 | 1 | 1 | 1 | 1 | 97,28 | 102,21 |
| Tyrosine Phosphatase | 1 | 0 | 1 | 1 | 0 | 0,00 | 1,16 |
| **Immunity total** | **100** | **66** | **88** | **96** | **70** | **15405,34** | **27765,52** |
