## Supplementary material for "Analysis of the testicle’s transcriptome of the Chagas disease vector *Rhodnius prolixus*": S4 Table

### S4 Table: Summary of secreted gene transcripts by functional protein families.

| Secreted proteins functional classes | # of transcripts | Expressed in Testes | Expressed in other tissues | Mapped to genome | Mapped to proteome | RPKM Testes | RPKM other tissues |
|---|---|---|---|---|---|---|---|
| 24 kda salivary protein | 1 | 1 | 1 | 1 | 0 | 139,62 | 10,31 |
| 47 kda salivary protein | 1 | 0 | 1 | 1 | 1 | 0,00 | 32,20 |
| Anterior gradient (agr) family | 2 | 2 | 0 | 2 | 2 | 468,03 | 0,00 |
| Antigen 59 | 1 | 1 | 1 | 1 | 1 | 36,29 | 35,03 |
| Apyrase | 1 | 0 | 1 | 1 | 1 | 0,00 | 3,32 |
| Basic tail | 2 | 1 | 2 | 2 | 0 | 101,65 | 58,78 |
| Cystatin | 1 | 1 | 1 | 1 | 1 | 125,63 | 664,09 |
| Endopeptidase | 1 | 1 | 1 | 0 | 1 | 32,16 | 19,08 |
| Ferritin | 6 | 4 | 6 | 4 | 4 | 261,80 | 343,43 |
| Fringe-like | 2 | 0 | 2 | 2 | 0 | 0,00 | 15,19 |
| Glutathione peroxidase | 1 | 1 | 1 | 0 | 1 | 28,99 | 43,01 |
| Haemolymph JHBP | 33 | 22 | 27 | 23 | 24 | 1552,45 | 1873,30 |
| Hyaluronidase | 3 | 3 | 2 | 3 | 1 | 24,75 | 4,96 |
| Inositol polyphosphate 5-phosphatase | 2 | 0 | 2 | 2 | 2 | 0,00 | 31,40 |
| Insect pheromone-binding family | 15 | 10 | 11 | 14 | 11 | 987,47 | 334,60 |
| Kazal type serpin | 11 | 10 | 5 | 8 | 5 | 1607,62 | 1727,45 |
| Lipocalin | 27 | 19 | 19 | 16 | 12 | 13307,01 | 8677,94 |
| Mucin | 1 | 1 | 0 | 1 | 1 | 28,74 | 0,00 |
| Mys precursor | 6 | 6 | 2 | 6 | 5 | 438,27 | 133,78 |
| Nitric oxide synthase | 2 | 1 | 2 | 2 | 2 | 9,82 | 15,88 |
| Nitrophorin | 6 | 6 | 4 | 5 | 4 | 300,38 | 88,61 |
| Odorant binding protein | 13 | 10 | 9 | 10 | 8 | 571,37 | 295,50 |
| Other secreted | 18 | 15 | 17 | 14 | 12 | 812,84 | 172,60 |
| Peptidase inhibitor | 2 | 1 | 2 | 2 | 1 | 13,74 | 13,30 |
| Platelet aggregation inhibitor | 1 | 0 | 1 | 0 | 0 | 0,00 | 20,27 |
| Protease inhibitors | 3 | 3 | 3 | 2 | 0 | 97,52 | 101,24 |
| Salivary odorant binding protein | 1 | 1 | 1 | 1 | 0 | 64,46 | 1,63 |
| Salivary phospholipase | 1 | 0 | 1 | 1 | 1 | 0,00 | 4,36 |
| Salivary secreted peptide | 19 | 14 | 19 | 18 | 10 | 1430,51 | 1508,12 |
| Scp | 4 | 3 | 4 | 4 | 3 | 192,04 | 107,28 |
| Serine protease inhibitor | 36 | 26 | 36 | 30 | 12 | 14240,83 | 11536,50 |
| Thrombin inhibitor rhodniin chain r | 6 | 6 | 6 | 3 | 0 | 458,82 | 595,99 |
| Triabin | 1 | 0 | 1 | 1 | 1 | 0,00 | 7,37 |
| Trypsin | 2 | 1 | 2 | 2 | 1 | 5,02 | 6,43 |
| Trypsin inhibitor heavy chain | 5 | 3 | 5 | 4 | 4 | 28,19 | 42,06 |
| Tryptophan rich secreted protein | 3 | 0 | 3 | 3 | 0 | 0,00 | 15,06 |
| Ubiquitin | 1 | 0 | 1 | 1 | 1 | 0,00 | 6,74 |
| Uncharacterized conserved protein | 2 | 1 | 1 | 2 | 0 | 38,85 | 6,05 |
| Unknown secreted | 14 | 8 | 11 | 14 | 6 | 331,86 | 222,87 |
| **Secreted total** | **257** | **182** | **214** | **207** | **139** | **37736,74** | **28775,75** |
