## Supplementary material for "Analysis of the testicle’s transcriptome of the Chagas disease vector *Rhodnius prolixus*": S5 Table

### S5 Table: Transcriptome mapping to proteome summary

| Alignment type | Unique proteins matching to trascriptome | Proteins with match to multiple transcripts | Total of transcripts hits to proteome |
|---|---|---|---|
| Without filter | 8,465 | 6,235 | 14,700 |
| ID ≥ 98 | 7,126 | 2,185 | 9,311 |
| ID ≥ 95 | 7,774 | 2,843 | 10,617 |
| ID ≥ 90 | 8,227 | 3,356 | 11,583 |
| ID ≥ 80 | 8,585 | 4,056 | 12,641 |
| ID ≥ 50 | 8,905 | 5,150 | 14,055 |
