## Supplementary material for "Analysis of the testicle’s transcriptome of the Chagas disease vector *Rhodnius prolixus*": S6 Table

**S6 Table: number of transcripts, by functional groups, that mapped to genome with 100% nucleotide identity but failed to map to proteome with >98% amino acid identity**

| Functional group | Count |
|---|---|
| **Housekeeping total** | **983** |
| Cytoskelletal | 97 |
| Detoxification | 9 |
| Detoxification/ox | 12 |
| Extransporteracellular matransporterix | 39 |
| Metabolism/aa | 7 |
| Metabolism/carb | 25 |
| Metabolism/energy | 22 |
| Metabolism/int | 9 |
| Metabolism/lipd | 49 |
| Metabolism/nuclear | 20 |
| Nuclear function | 58 |
| Nuclear membrane | 5 |
| Protein machinery | 24 |
| Protein machineryein export | 47 |
| Protein machineryein metabolism | 44 |
| Protein metabolism/protease | 29 |
| Protein synthesis | 42 |
| Signal transduction | 204 |
| Signal transduction/apoptosis | 13 |
| Transcription factor | 35 |
| Transcription machinery | 129 |
| Transporter | 64 |
| **Immunity total** | **9** |
| Defensin | 1 |
| Lactalbumin | 1 |
| Other immunity | 3 |
| Ras gtpase activating protein | 1 |
| Rhomboid | 1 |
| Suppressor of cytokine | 1 |
| Tumor necrosis factor | 1 |
| **Secreted total** | **42** |
| Basic tail | 2 |
| Ferritin | 1 |
| Fringe-like | 1 |
| Haemolymph JHBP | 4 |
| Insect pheromone-binding family | 1 |
| Kazal type serpin | 2 |
| Lipocalin | 5 |
| Odorant binding protein | 1 |
| Other secreted | 1 |
| Peptidase inhibitor | 1 |
| Protease inhibitors | 2 |
| Salivary odorant binding protein | 1 |
| Salivary secreted peptide | 4 |
| Scp | 1 |
| Serine protease inhibitor | 11 |
| Thrombin inhibitor rhodniin chain r | 1 |
| Tryptophanrich secreted protein | 1 |
| Uncharacterized conserved protein | 1 |
| Unknown secreted | 1 |
| **Transposable element** | **23** |
| **Unknown conserved** | **317** |
| **Unknown** | **1922** |
| **Viral** | **1** |
| **Total** | **3297** |
